# *Theileria annulata* Infection Promotes p53 suppression, Genomic Instability and DNA deaminase APOBEC3H upregulation leading to cancer-like phenotype in host cells

**DOI:** 10.1101/2024.02.20.581323

**Authors:** Debabrata Dandesena, Akash Suresh, Roli Budhwar, Jeffrey Godwin, Sakshi Singh, Madhusmita Subudhi, Amruthanjali T, Sonti Roy, Vengatachala Moorthy A, Vasundhra Bhandari, Paresh Sharma

## Abstract

*Theileria annulata*-infected host leukocytes display cancer-like phenotypes, though the precise mechanism is yet to be fully understood. The occurrence of cancer-like phenotypes in *Theileria*-infected leukocytes may be attributed to various factors, including genomic instability and acquired mutations, a crucial trait that underpins the genetic foundation of cancer. This paper presents WGS data and bioinformatic analyses to reveal point mutations and large-scale alterations in six clinically relevant *T. annulata*-infected cell lines. We identified 7867 exon-linked somatic mutations common to all cell lines, and cancer association analysis showed significant accumulation in oncogenes (FLT4, NOTCH2, MAP3K1, DAXX, FCGR2B, ROS1) and tumor suppressor genes (BARD1, KMT2C, GRIN2A, BAP1) implicated in established critical cancer processes. We demonstrated that a crizotinib-induced blockade of the ROS1 oncogenic protein, which harbored the most mutations, led to the death of infected leukocytes. This is consistent with the significant role of ROS1 in parasite-induced leukocyte transformation. In addition, we found somatic mutations in genes involved in genome instability and the DDR pathway. Our findings support the notion that ROS1 and Nutulin 3a are valid targets for intervention, and the suppression of TP53, a crucial tumor suppressor gene, may play a significant role in cell immortalization. We also show that upon infection with the parasite, bovine cells upregulate the expression of APOBEC3H, a DNA mutator likely responsible for the detected mutations. Our study highlights how *T. annulata* transforms leukocytes to gain selective advantage via mutation, and our observations could steer future research towards a mechanistic understanding of disease pathogenesis.

## Introduction

Parasites in the phylum Apicomplexa cause significant morbidity and mortality in humans and animals worldwide. Among these, *Theileria annulata* and *Theileria* parva are the only parasites shown to trigger host cell transformation, a process with many of the hallmarks of cancer (Tretina *et al*., 2015, Rchiad Z., 2020; Cheeseman K *et al*., 2021; Li Z *et al*., 2021; Villares *et al*., 2022). Sporozoites of *T. annulata* are capable of infiltrating lymphocytes and macrophages, leading to the development of macroschizonts which can induce a malignancy in the host cells (Tretina *et al*., 2015). The development of cancer-like characteristics in *Theileria*-infected leukocytes is associated with epigenetic modifications and changes in host gene expression (Dobbelaere and Rottenberg, 2003; Cock-Rada *et al*., 2012; Kinnaird *et al*., 2013; Tretina *et al*., 2015; Robert MW *et al*., 2016; Woods *et al*., 2021). However, the precise mechanism through which parasite-infected leukocytes develop the cancer-like features remains unknown. Since *Theileria*-infected leukocytes share many characteristics with cancer, a genetic disease, it’s plausible that somatic mutations in host genes might offer a selective advantage in suppressing apoptosis and inducing uncontrolled host cell proliferation (Tretina *et al*., 2015).

Among the numerous potential causes of cancer, pathogen-induced DNA damage contributes to genomic instability and mutations in about 20% of cancers (Zur Hausen, 2009; de Martel *et al*., 2012; Matthew & Jonathan., 2014). To survive and thrive, pathogens may directly or indirectly cause host cell damage that impairs their genomic integrity (Matthew & Jonathan., 2014). Loss of integrity can lead to genomic instability, a hallmark of cancer cells mainly driven by mutations in DNA repair genes, or faults in DNA replication genes (Yao & Dai, 2014; Huang & Zhou,2021). Both cancer and Theileria-transformed leukocytes are associated with uncontrolled cell growth, which may lead to genomic instability and an increased propensity for mutation acquisition. Infection of leukocytes by *Theileria* parasites leads to upregulated expression of some host genes such as MDM2 (mouse double minute 2 homologs; p53 negative regulator) and SMYD3 (SET and MYND domain containing 3), which may induce genomic instability in infected cells, although no direct relationship has been demonstrated (Baylin & Herman; 2000; Robertson., 2001; Haller *et al*., 2010; Tili *et al*., 2011; Cock-Rada *et al*., 2012; Hayashida *et al*., 2013; Matthew & Jonathan., 2014). Besides *Theileria*, *Cryptosporidium*, another apicomplexan parasite, is also recognized for inducing cancer-like symptoms in experimental models through disruption of host DNA integrity (Matthew & Jonathan., 2014; Tretina *et al*., 2015).

Because cancer is a genetic disease, mutations are extensively studied for any connection to cancer, but their potential significance in *Theileria*-infected leukocytes remains unclear (Watson *et al*., 2013; Marco A *et al*., 2022). Cancer-like characteristics, such as genomic instability and acquired mutations, have not yet been explored in *Theileria*-transformed leukocytes. Genomic instability and mutation in *Theileria*-infected leukocytes should be examined to see whether infection and resulting cellular transformation provoke instability and/or changes to the host genome, and if so, does parasite infection cause cancer to host cells?

Next-generation sequencing has made it possible to systematically discover the mutational spectra of various cancers that may arise from inherited mutations, environmental causes, or faulty DNA replication (Tomasetti *et al*., 2017). Here, we utilized WGS to investigate the effect of *T. annulata* infection on host genome integrity and the function of somatic mutations in determining whether there is a genetic connection to the cancer-like phenotype of infected and transformed leukocytes. We postulated that *Theileria*-infected leukocytes might gain a selective advantage due to mutations in oncogenes/ tumor suppressor genes (TSGs), revealing previously unknown aspects of parasite biology. We identify linkages that enhance our knowledge of the *Theileria* parasite’s adaptation to its host leukocyte, a significant portion of which has not been previously investigated. Identifying these traits opens the path for creating and using cancer therapeutics to treat tropical theileriosis caused by *T. annulata* and a greater understanding of the molecular connections underlying the resulting cancer-like phenotypes.

## Result

### Whole genome characterization of somatic mutations in lymphoproliferative cells infected with *T. annulata* parasites

To study the impact of *T. annulata* infection on host genome integrity, we conducted WGS analysis on six transformed leukocyte cell lines derived from PBMCs of clinically diseased cattle *(Bos taurus*) in India. Confirmation of parasites in all cell lines was made by utilizing PCR and IFA, which were directed towards TASP, a gene exclusive to *T. annulata* parasites. An IFA image demonstrating the presence of *T. annulata* in these infected cell lines is shown (Supplemental_Fig1.pdf). The six cell lines have been previously characterized using microsatellite markers and genotype-based sequencing analysis (Roy *et al*., 2019). We analyzed the sequenced samples using the *Bos taurus* genome (ARS UCD 1.2), as a reference to identify changes in cancer-related genes. Table 1 shows the number of reads per sample, and a flowchart of the overall WGS statistics plus workflow is shown (Supplemental_Fig2.f). Variants were evaluated only if they met the following criteria: a phred quality score of at least 30, an allele frequency of at least 8%, and a significance level of P <0.05. Variants with a variant allele frequency of >60% were called homozygous. Furthermore, we excluded non-coding data concentrating on coding regions to identify synonymous, and non-synonymous (missense) mutations. In recent years it has become clear that synonymous mutations may also contribute to cancer and are no longer considered neutral (Sharma *et al*., 2019; Westcott *et al*., 2015; Shen *et al*., 2022). Each sample had an average of 98,000 somatic mutations (SNPs) and 3400 indels in coding regions (Fig: 1A,1B). We next looked for alterations common to all samples to identify genes that might play a role in inducing the cancer-like phenotype of *Theileria*-infected leukocytes. A total of 7867 shared mutations were split between non-synonymous (n=3590) and synonymous (n=4262) variants, representing 3,580 genes (Supplemental_Table1.xls). There were 1136 cases of homozygous and 6731 cases of heterozygosity. Besides the single gene variations, we also detected 412 common indels in genes across the WGS data sets (Fig: 1B)

**Figure 1:**
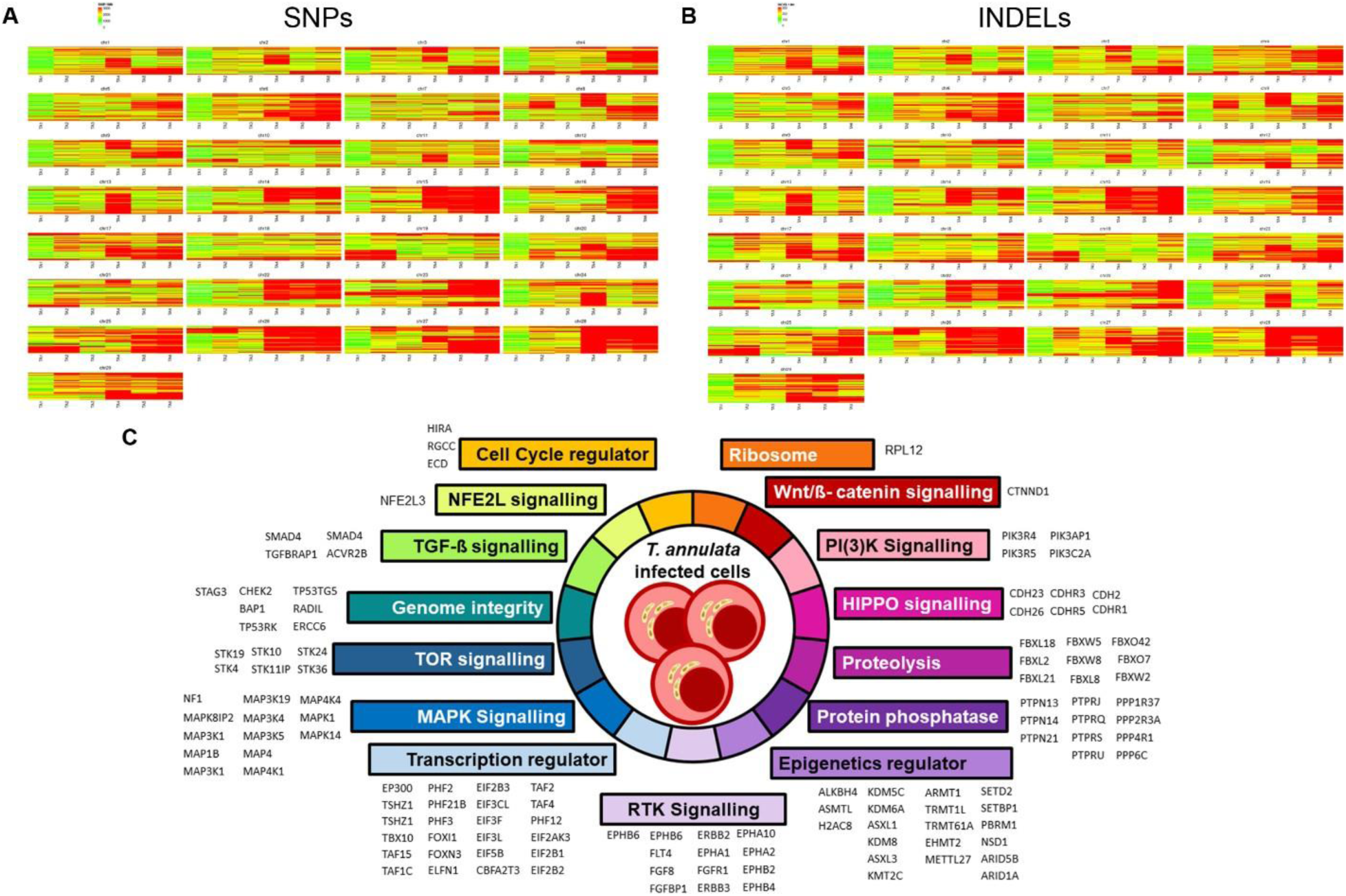
Chromosome-wise distribution of SNPs, INDELs, and significantly altered pathways in *T. annulata* infected samples. (A) The heat map represents the chromosome-wise distribution of all the SNPs across 29 chromosomes for 6 WGS samples used in the study. (B) Heat map representing the chromosome-wise distribution of all the Indels for 6 WGS samples used in study (C) Cancer enrichment analysis using TCGA database showing major signaling pathways altered across our samples based on the gene mutations. The figure shows pathways and related genes mutated across 6 *Theileria* infected cells identified using WGS analysis.

Furthermore, we used the TCGA and COSMIC databases to identify 127 genes that may be implicated in the cancer-like phenotype of *Theileria*-infected leukocytes. Comparative analysis identified significant mutations in the genes known to have an important role in cancer. The TCGA dataset provides a comprehensive list of genes that are known to participate in various cellular functions across multiple types of cancers (Kandoth *et al*., 2013). By analyzing our own data sets, we have identified 127 distinct significantly mutated genes (SMGs), indicating the potential involvement of these cellular and enzymatic mechanisms in *Theileria*-induced leukocyte transformation. As per TCGA dataset, these SMGs play crucial roles in essential cellular processes associated with cancer, including cell cycle regulation, mitogen-activated protein kinase (MAPK) signaling, phosphatidylinositol-3-OH kinase (PI(3)K) signaling, Wnt/ß-catenin signaling, transcription factors/regulators, epigenetic regulation, maintenance of genome integrity, and ubiquitin-mediated proteolysis. (Fig: 1C).

Next, using the COSMIC database, we searched for mutations in cancer hallmark genes involved in oncogenic processes such as proliferation, growth suppression, replicative cell immortality, invasion and metastasis, genomic instability and mutations, evasion of programmed cell death, and changes in cellular energy metabolism. Except for genomic instability and mutations, these cancer-like characteristics are documented in the context of the *Theileria*-infected leukocytes (Tretina *et al*., 2015). We discovered 121 mutated genes common to all 6 samples, 89 of which were ranked as “tier-1” and 32 as “tier-2” in the COSMIC database. Since mutations in “tier-1” genes have been directly shown to promote cancer, we next identified 22 genes with non-synonymous mutations (Supplemental_Table1.xls). In addition to the common missense mutations in the 22 COSMIC tier-1 genes, each gene also possessed unique alterations that may be functionally significant for its encoded activity. The bar graph depicts the mutational load of COSMIC tier-1 genes (n=22) in *Theileria*-infected leukocytes (Fig: 2A). Twelve of these tier-1 genes are also cancer hallmark genes, as mutations in them have been demonstrated to have a direct role in cancer progression: FLT4, NOTCH2, MAP3K1, DAXX, BARD1, KMT2C, GRIN2A, BAP1, FCGR2B, ROS1, SLC34A2, and NOTCH1 (Fig: 2B). Using the SIFT score, we examined the influence of non-synonymous mutations on the cosmic listed genes in our datasets. We discovered mutations in KMT2C, TSC2, ROS1, HOOK3, LZTR1, CHEK2, TAF15, and GNA11 that might impact their function. KMT2C and ROS1 were discovered to have the highest number of mutations in our samples. Figures 2C & 2D show a lollipop plot for KMT2C and ROS1, illustrating the distribution of variants and their effect on the genes. Since we observed the highest number of mutations in ROS1, *Theileria*-infected leukocytes were challenged with crizotinib, a tyrosine kinase inhibitor approved by the US Food and Drug Administration (FDA) for the treatment of certain types of advanced non-small cell lung cancer (NSCLC) harboring ROS1 mutations (Shaw *et al*., 2014). By blocking the activity of the ROS1 protein, crizotinib inhibited growth (IC50= 8.89µM) of *Theileria*-transformed leukocyte cell lines (n=3), highlighting its potential importance in establishing their cancer-like phenotype (Supplemental_Fig3).To assess the impact of mutations on specific candidate tumor suppressor genes (BARD1, KMT2C, GRIN2A, SLC34A2, NOTCH1, and BAP1), we conducted q-PCR analysis on cell lines infected with *Theileria*, using healthy PBMCs as a control. Our findings revealed a significant decrease in the expression of all the examined tumor suppressor genes in the parasite-infected lines. These mutations may lead to the absence or impaired functioning of the proteins encoded by these genes, potentially resulting in uncontrolled cell division and contributing to the development of cancer-like phenotypes in the parasite-infected cells (Fig 2E).

**Figure 2:**
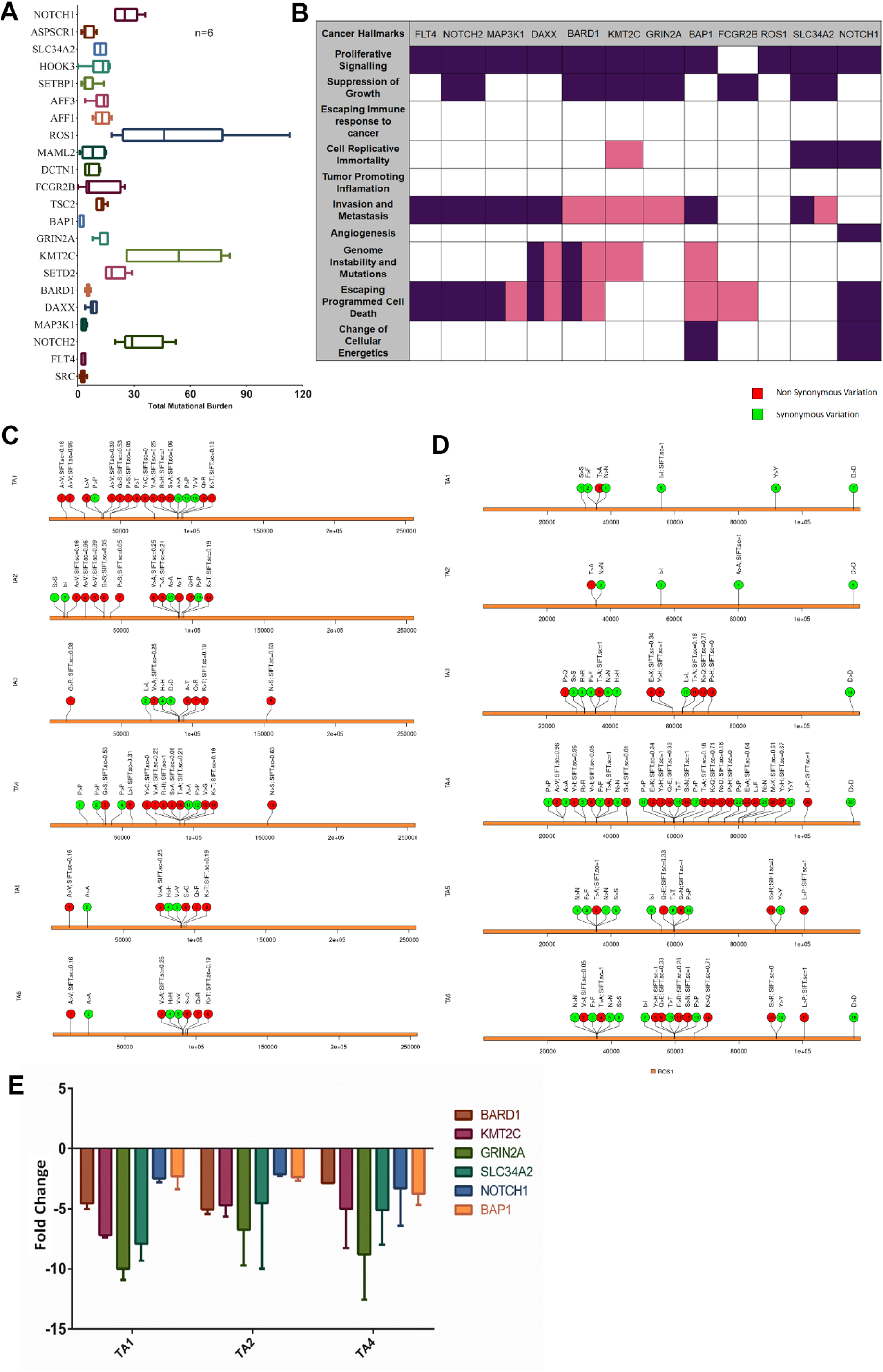
Mutation in the cancer hallmark genes and their functional relevance in *T. annulata* infected samples. (A) The graph shows the total mutational burden of the 22 tier 1 gene (COSMIC) identified in our data using WGS analysis. Gene’s mutational burden was calculated individually in each sample and then averaged for the six samples. Both synonymous and non-synonymous mutations were calculated for all six samples and represented in the form of a box plot as the total mutational burden. Error bars represent the mean with standard deviation (SD). (B) Based on the COSMIC database, the figure shows the list of cancer hallmark genes and their process linked. These 12 cancer hallmark genes are found to have common non-synonymous mutations throughout the six samples in our data. These genes are shown to have roles as tumor activators or suppressors marked with purple and pink colors respectively. (C) & (D) Lollipop images representing the distributionof mutations along with the SIFT score of KMT2C and ROS1 across six *Theileria*-infected cells. (E) q-PCR based gene expression levels of tumor suppressing genes in *T. annulata* cells w.r.t uninfected healthy PBMCs.

Alterations in genes associated with the genomic instability pathway, such as DAXX, BARD1, KMT2C, and BAP1, may explain the increased frequency of mutations in *Theileria*-infected cells. A defect in the DNA repair pathways increases the risk of cancer and genomic instability (Abbas *et al*., 2013). We discovered common mutations in 58 genes involved in important DNA repair pathways across our datasets (Supplemental_Table2.pdf). The major DDR- related genes and pathways that have been mutated are homologous recombination (MUS81, XRCC2, PALB2, RPA2, EME1), non-homologous end joining (PRKDC, MAD2L2), miss match repair (PMS1), nucleotide excision repair (GTF2H4, LIG1, UVSSA, ERCC6, GTF2H3, CCNH), base excision repair (NEIL2), fanconi anaemia repair (FANCD2, FANCM, FANCC), DNA polymerase (POLI, POLN, POLG, POLD1, POLE), ubiquitin and modification (RNF8), repair of DNA-protein crosslinks (TDP1), and other conserved DDR pathways proteins like CLK2, CHEK2, TOPBP1and MDC1 (Fig: 3A). CHEK2, a TSG gene generally engaged in DNA repair, had both shared and distinct synonymous and non-synonymous mutations across the samples (Fig: 3B). The detected mutations in the CHEK2 gene have a deleterious effect (SIFT<0.05) on the gene’s normal function, indicating a likely DNA repair pathway defect in *Theileria* infected cells.

**Figure 3:**
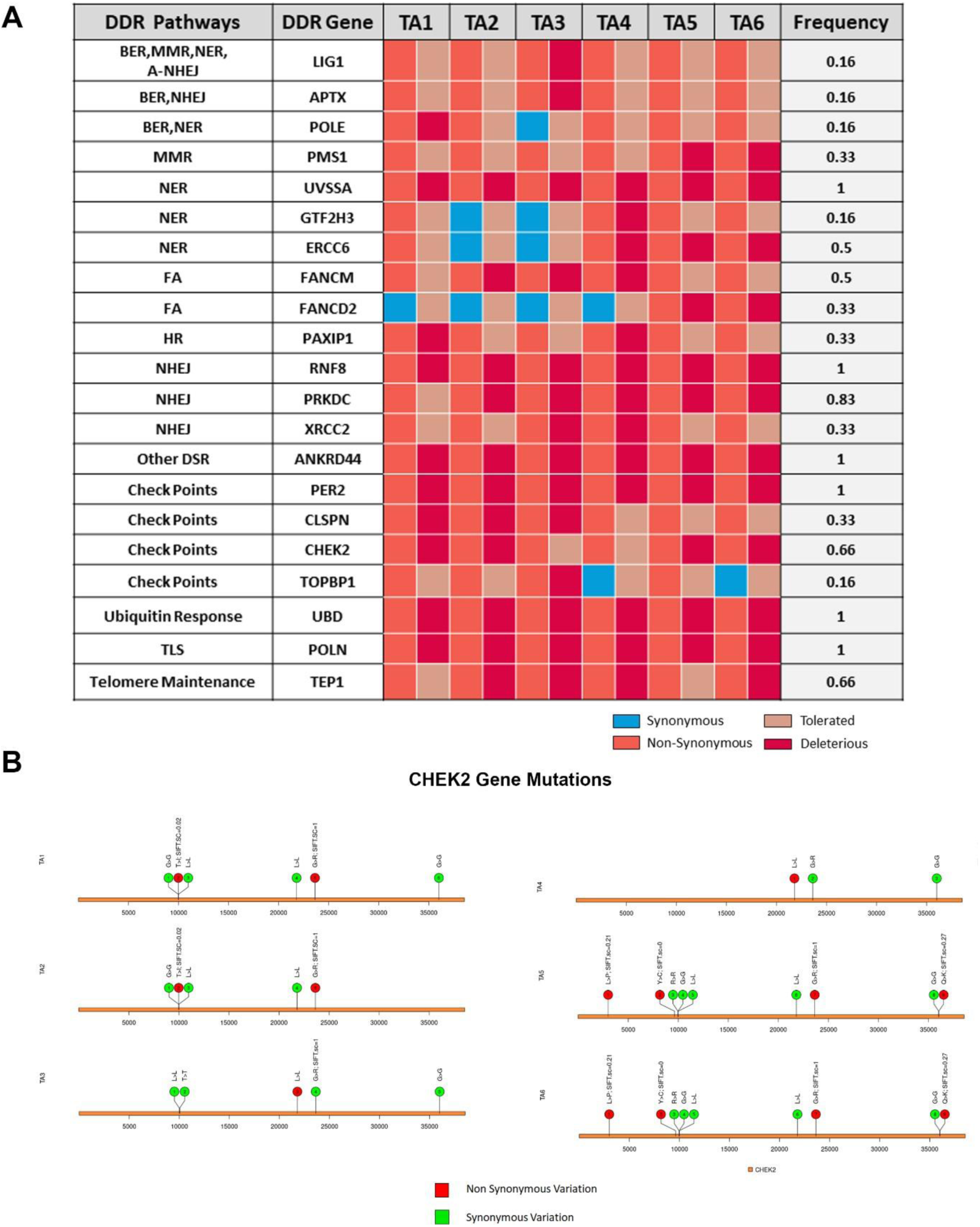
Mutational spectrum of DDR genes in the *T. annulata* infected samples. (A) The figure shows major DDR pathway-related genes altered in our samples and the frequency of their alteration in the six datasets. Synonymous mutations are marked with blue and non-synonymous with orange color. The impact of the mutations was marked with another two sets of colors, light orange represents the tolerated mutation, and red represents the deleterious mutations predicted from the SIFT score. Samples lacking any mutation for the genes are marked with grey color. *Represent the presence of missense variation. The frequency of each gene is represented at the end. (B) Lollipop image showing the mutations of the CHEK2 gene for all six samples along with the SIFT score. All six samples have multiple mutations representing non-synonymous(red) and synonymous(green). Four out of the six samples had deleterious mutations, marked by the SIFT score.

In addition to cancer-related gene mutations, we found common homozygous non-synonymous alterations in epigenetic regulators such as KMT2C, ARMT1, TRMT1L, TRMT61A, EHMT2, METTL27, ALKBH4, ASMTL, H2AC8 and EHMT2 (Supplemental_Table1.xls). Since epigenome regulator mutations are possible therapeutic targets in cancer, they might be crucial in the life cycle of *T. annulata* and beneficial in developing therapies against the parasite (Plass *et al*., 2013).

### DNA mismatch repair (MMR) related signatures leads to Microsatellite instability (MSI) in *Theileria*-infected cells

WGS has advanced cancer genetics to the point that cancer-type-specific mutational signatures have been discovered (Degasperi *et al*., 2022). These mutational fingerprints represent endogenous and exogenous cancer factors (Alexandrov *et al*., 2015; Islam *et al*., 2022).

Multiple mutational mechanisms cause somatic mutations in cancer genomes, each leaving a mark. Mutational signature analysis may assist in determining if there is a relationship between observed mutations and genomic instabilities, or defective DNA repair pathways. Since mutational signatures for most cancer types are readily accessible, this analysis would also help us determine whether *Theileria*-transformed leukocytes and human cancer share a common signature. Based on the orientation of mutations, all point substitutions were classified into six types (C>A, C>G, C>T, T>A, T>C, T>G) using SomaticSignatures (Fig: 4A, B, C). The proportions of all six mutation groups were comparable across samples, with C>T and T>C being the most prevalent mutation types based on the sequencing data (Supplemental_Table3.xls). Using deconstructSigs we next established if any mutational signatures are associated with specific mutation subtypes. It revealed signatures characteristic of SBS1A, SBS20, and SBS12 in all six infected leukocyte lines (Fig: 4D). SBS1A (associated with deamination of 5-methyl cytosine) signatures are ubiquitous among different cancer types and present in most cancer cell lines (Alexandrovcf *et al*., 2015). SBS20 is associated with concurrent POLD1 mutations, defective DNA mismatch repair (MMR), and microsatellite instability (MSI) (Aleksandrov *et al*., 2018). POLD1 mutation was present across all of 6 of the infected leukocyte cell lines, validating the results of the mutational signature (Supplemental_Table1.xls). Mutations in DNA mismatch repair genes (MMR) induce microsatellite instability (MSI), a hypermutable trait that increases cancer risk (Pecina-Slaus *et al*., 2020). Four genes MLH1, MSH2, MSH6, and PMS2 govern the fate of the MMR pathway. We next asked if any mutations were present in the four MMR genes in *Theileria*-infected leukocytes. Both MSH2 and PMS2 were mutated in all six *Theileria*-infected leukocyte cell lines, while MSH6 was mutated in all but one line. In addition, mutations were also found in additional MMR genes (MSH3, MSH4, MSH5, PMS1, HFM1, MSX2) in five of the *Theileria*-infected leukocyte cell lines (Supplemental_Table3.xls). As MMR may result in microsatellite instability (MSI), we next examined the *Theileria*-infected leukocyte cell lines for the presence of MSI (Supplemental_Table4.xls). Using the WGS datasets, we measured the lengths of specific DNA microsatellites present in *Theileria*-infected leukocytes (Fig:4 E). MSI was detected in all 6 infected leukocyte cell lines, with a mean MSI index of 24.19 ± 3.38 per sample (Fig:4 F).

**Figure 4:**
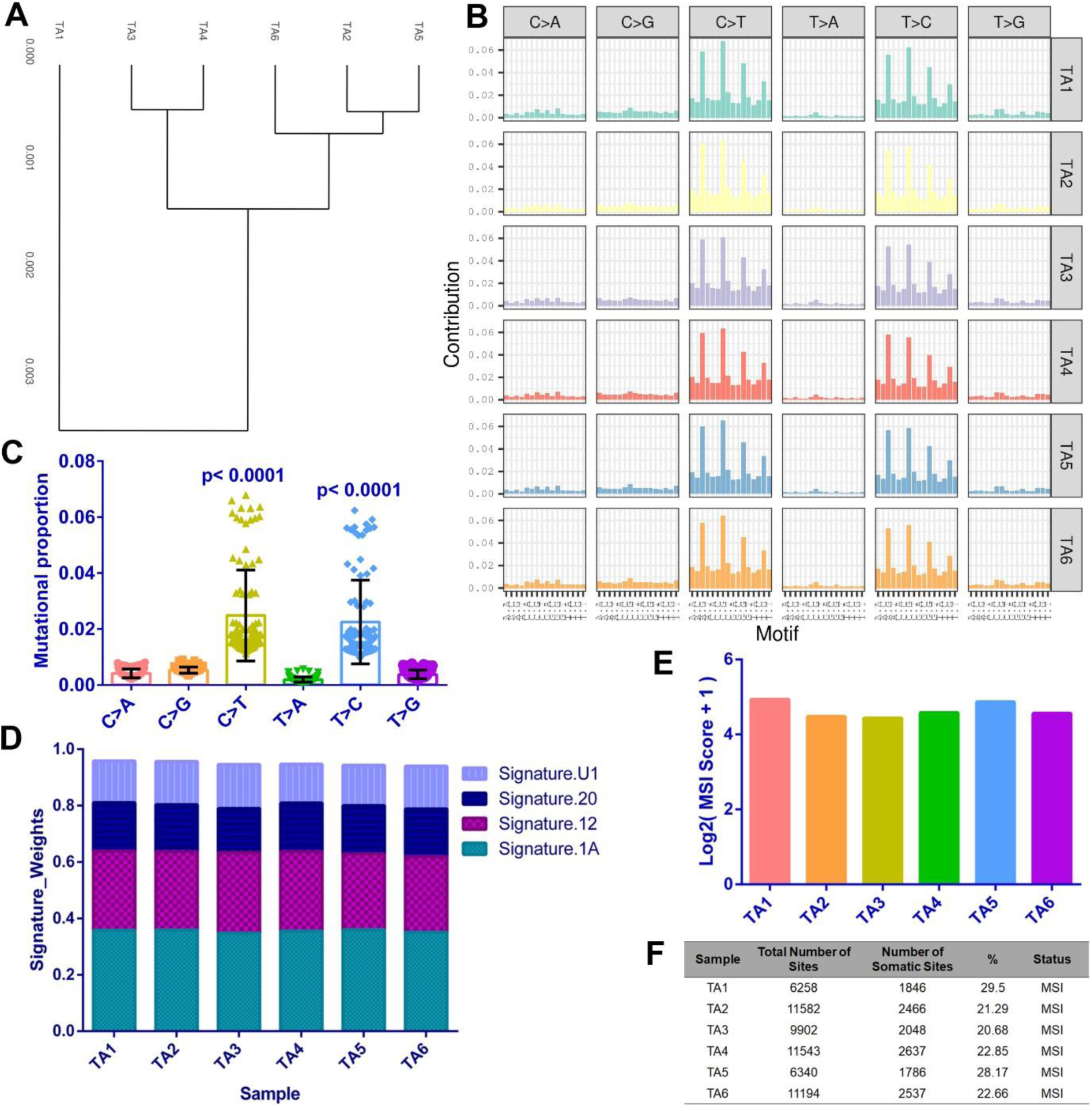
Divergent mutational features and MSI status of *T. annulata* infected bovine lymphocytes compared to the reference. (A) Dendrogram represents the hierarchical clustering of the mutational spectrum across six *T. annulata* infected samples. (B) Contribution of six mutational subtypes (C>A, C>G, C>T, T>A, T>C, T>G) represented in the form of a bar chart for all six different datasets (TA1-TA6), marked with different color codes. (C) C>T and T>C substitutions are significantly elevated across all six samples compared to the other four. Mutational proportion data of all the 16 possible tri codons for each substitution for all six samples have been represented as a scattered plot. The error bars reflect mean and standard deviations. One-way ANOVA analysis was carried out to perform multiple comparisons among all six single-base substitutions. The graph represents the frequency of each substitution for all six samples. Tukey’s multiple comparisons test was used and the p-value was calculated **** (p-value<0.0001). (D) **Mutational signature present in all *T. annulata* infected samples**: We reconstructed the proportion of mutational signatures of each sample based ona predefined mutational spectrum of 30 COSMIC signatures. The four identified signatures in the infected samples are shown. (E) The bar graph displays the MSI score of all six samples in the Log2(MSI Score+1) scale. (F) The table represents the percentage of MSI sites along with the MSI status of all six samples.

### Copy number variation (CNV) analysis identified alterations in cancer-related genes in the *Theileria*-infected cells

CNVs influence more of the genome than other somatic mutation in cancer cells, activating oncogenes and inactivating TSGs (Shao *et al*., 2019). Therefore, we asked whether there were any CNVs associated with the cancer-like phenotype of *Theileria*-infected and transformed leukocytes. We exploited CNVcaller to identify the CNVR (copy number variation regions) across the six *Theileria*-infected leukocyte cell lines (n=6) (Fig: 5A, 5B, 5C, 5D). We detected 66,171 CNVR to be common to the six *Theileria*-infected leukocyte cell lines associated with both gene amplifications (61,121) and deletions (5050) (Supplemental_Table5.xls).

**Figure 5:**
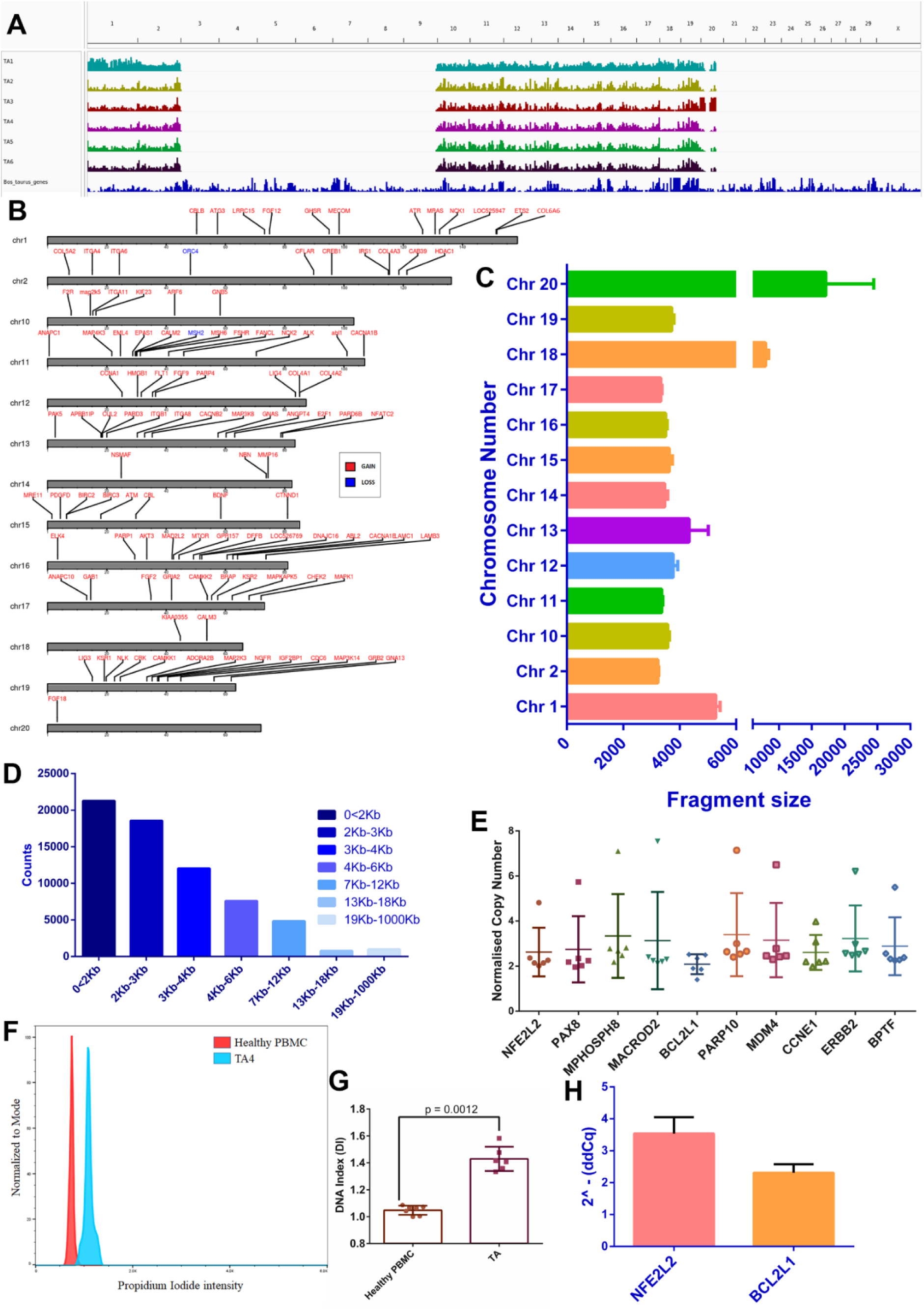
Analysis of CNVs of six *T. annulata* infected samples. (A) The graph shows a chromosomal bed representing the distribution of CNVs across six samples in the 29 chromosomes using *B. taurus* genome as a reference. (B) The figure shows the chromosome-wise distribution of common genes with a gain or loss of copy number marked by blue and red across the six samples. (C) Bar graph representing the chromosome-wise distribution of fragment size (both amplification and deletion). The error bar represents the mean with SD. (D) The fragment sizes were divided into different subtypes based on the length. The absolute count for each subtype was plotted in the form of a bar graph. (E) Scatter plot showing the normalized copy number for the genes that were found to be significantly altered across many cancer types. (F) Histogram representing the flowcytometry-based DNA content in *T. annulata* infected cells w.r.t healthy PBMCs. (G) Representation of DNA index in the form of the bar graph. (H) The q-PCR-based relative gene expression analysisof two genes, NFE2L2 and BCL2L1, is shown using HPRT as housekeeping control. The fold change was calculated using 2^^-ΔΔCT^, and the error bar represents the standard error mean.

Next, we searched for CNVs in the cancer-related genes using the 2013-08-2016 TCGA Pan-cancer dataset. The analysis unveiled amplification of 10 genes that are frequently reported as significantly altered in various types of cancers, within the *Theileria*-infected cell lines (Fig: 5E). The genes were: NFE2L2, MPHOSPH8, MACROD2, BCL2L1, PARP10, MDM4, CCNE1, ERBB2, PAX8, and BPTF. DNA gains and losses are common in cancer and have been shown to induce aneuploidy and chromosomal instability (Ippolito *et al*., 2021, Steele *et al*., 2022). Considering that healthy cells typically cannot withstand aneuploidy, our subsequent inquiry focused on determining if leukocytes infected with *Theileria* exhibit any signs of altered DNA content. We utilized DNA flow cytometry to evaluate the ploidy level in *T. annulata*-infected leukocytes (N=2) using healthy cattle leukocytes, as a reference for comparing DNA content. Interestingly, the DNA index of *T. annulata*-infected leukocytes exceeded 1.1, indicating aneuploidy. The initial peak on the left corresponds to diploid cells that are considered normal. The subsequent prominent peak indicates cells with an elevated DNA content and a DNA index of 1.58(Fig). These findings align with the WGS data, where we analyzed the 6 datasets for copy number variations that could potentially contribute to aneuploidy.

Different malignancies and cancer cell lines all share a significant association between CNV and altered gene expression (Shao *et al*., 2019). For independent confirmation of the CNV discovery-based approach q-PCR analysis of two candidate genes BCL2L1, and NFE2L2 was performed. This revealed increased expression of the anti-apoptotic gene BCL2L1 and the transcription factor NFE2L2, which both showed amplification in the CNV dataset (Fig: 5F). Enhanced mRNA expression of the BCL2L1 and the presence of a CNV in the gene might be responsible for the resistance to apoptosis phenotype observed in *Theileria*-infected and transformed leukocytes (Tretina *et.al*., 2015).

### Identification of large-scale Structural variations across *T. annulata* infected cells

Genomic deletions, duplications, and rearrangements may affect anything from a few kilobases to an entire chromosome, known as structural variation (SVs), which plays a criticalrole in the development of cancer (Bignell *et al., 2*007, Dixon *et al., 2*018, Li *et al., 2*020). Next,we wanted to discover similar genetic alterations in *Theileria*-infected cells since SVs play a vital role in changing gene expression and are one of the leading factors generating the cancer phenotype. We analyzed the genomes of the *Theileria*-infected cells using the algorithm Breakdancer to identify the four major somatic structural variants: deletions (DEL), inversions (INV), and translocations (CTX/ITX). This algorithm helped us to detect huge SVs (>1Kb), but for further analysis, we only took SVs with a confidence score of 99 (Supplemental_Table6.xls). Figure 6 A, B, C, D, E, F illustrates the SVs discovered in each of the six-cell lines using the CIRCOS plot. On average, 2427.3 ± 575.29 variations were detected across all samples. Out of which, 17.32 ± 6.53, 36.88 ± 10.81,15.84 ± 12.51, and 29.93 ± 9.03% belong to CTX, DEL, INV, and ITX, respectively (Fig: 6G). Sixteen genes had common SVs across all the *Theileria*-infected cells; these include SESTD1, ITPR2, ELMO1, IGF2BP2, HAT1, PPM1H, UBR4, DPP6, ZNF654, PDE5A, LNP1, GRID2, CGGBP1, RAP1A, CACNA1C, FABP2 (Fig: 6H). Our investigation of SVs led us to RAP1A, a protein belonging to the RAS oncogene family that controls signalling pathways affecting cell proliferation and adhesion and may play a role in tumor malignancy (Zhang *et al., 2*017).

**Figure 6:**
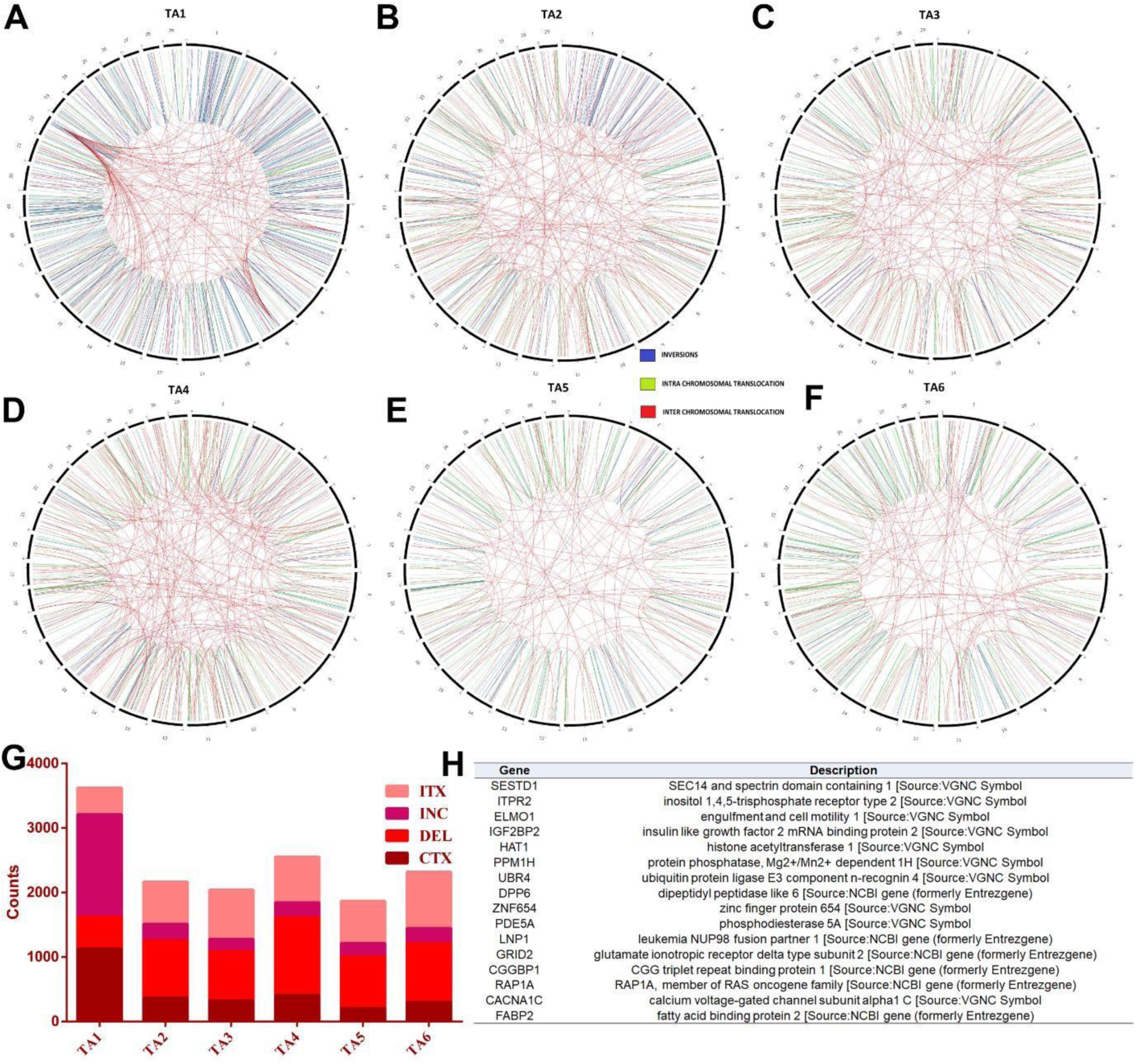
Distribution of structural variations across six *T. annulata* infected samples. (A-F) The chromosome-wise distribution of SVs like INV, ITX, and CTX is represented as a CIRCOS diagram for all six samples. (G) The bar graph shows the absolute count of four different SVs (INV, ITX, DEL, and CTX) detected across all six samples marked with different color codes. (H) The table shows the list of the genes and their function that have common SVs across six samples.

### Non-coding mutational analysis

Conventional efforts to find mutations that cause cancer have focused on decoding protein-coding genes. Non-coding mutations play a crucial role in human cancer, and WGS has found multiple cancer-causing changes in non-coding regulatory elements, including promoters, enhancers, intergenic regions, and 5′- and 3′-untranslated regions (5- and 3′-UTRs) (Kikutake *et al., 2*021; Sherman *et al., 2*022). Based on the analysis of non-coding somatic mutations, over 95% of the total mutations in *Theileria*-infected cells were in non-coding regions. Then, we searched for genes with somatic mutations in the non-coding regions of *Theileria*-infectedcells that may be associated with cancer. The non-coding analysis discovered mutations in the intergenic regions of the cancer-related genes TAL1, MCL1, MMP1, HGF, LMO1, SDHD, PAX5, MMP2, CASP8, NFKB1, BRAC1, MMP3, MMP7, CDKN1B, ERCC5, RAD51, RB1, BRCA2, APC, MSH6, PSMD8, CD59, CIAO1, SRSF5, and NDUFA6 to have homozygous mutations in our samples (Supplemental_Fig2.pdf). However, we found no common alterations in the non-coding genes across the samples.

#### TP53 mediates the induction of apoptosis in *Theileria*-infected leukocytes

The administration of buparvaquone (BPQ) leads to the death of *Theileria* parasites within host leukocytes. Without live parasites, host leukocytes lose their immortalized phenotype leading to activation and nuclear localization of the TP53 protein and, ultimately, leukocyte death (Haller *et al*., 2010). Therefore, we aimed to determine whether infected leukocyte immortalization is solely due to harboring live *Theileria* parasites, or if there are host factors that contribute to this cancer-like phenomenon. We tested the effects of Nutulin 3a, a TP53 gene activator, on Theileria-infected leukocytes to investigate the role of host. The administration of Nutulin 3a caused the activation of TP53, resulting in cell death much like the effects of BPQ treatment (Figure 7). This suggests that Nutulin 3a is able to directly activate TP53 by preventing the TP53-Mdm2 interaction, which leads to apoptosis of the infected leukocytes. This implies that while Theileria parasites are alive, TP53 is bound to Mdm2 to maintain survival of the infected leukocytes. TP53 has been identified as a key regulator in cancer, and its ability to induce programmed cell death is important for tumor suppression (Bieging *et al*., 2014). Our WGS findings, which show that there are no SNPs in TP53 within Theileria-transformed leukocytes, are validated by the promotion of apoptosis in infected leukocytes when TP53 is released from its binding to Mdm2 following treatment with Nutulin 3a.

**Figure 7:**
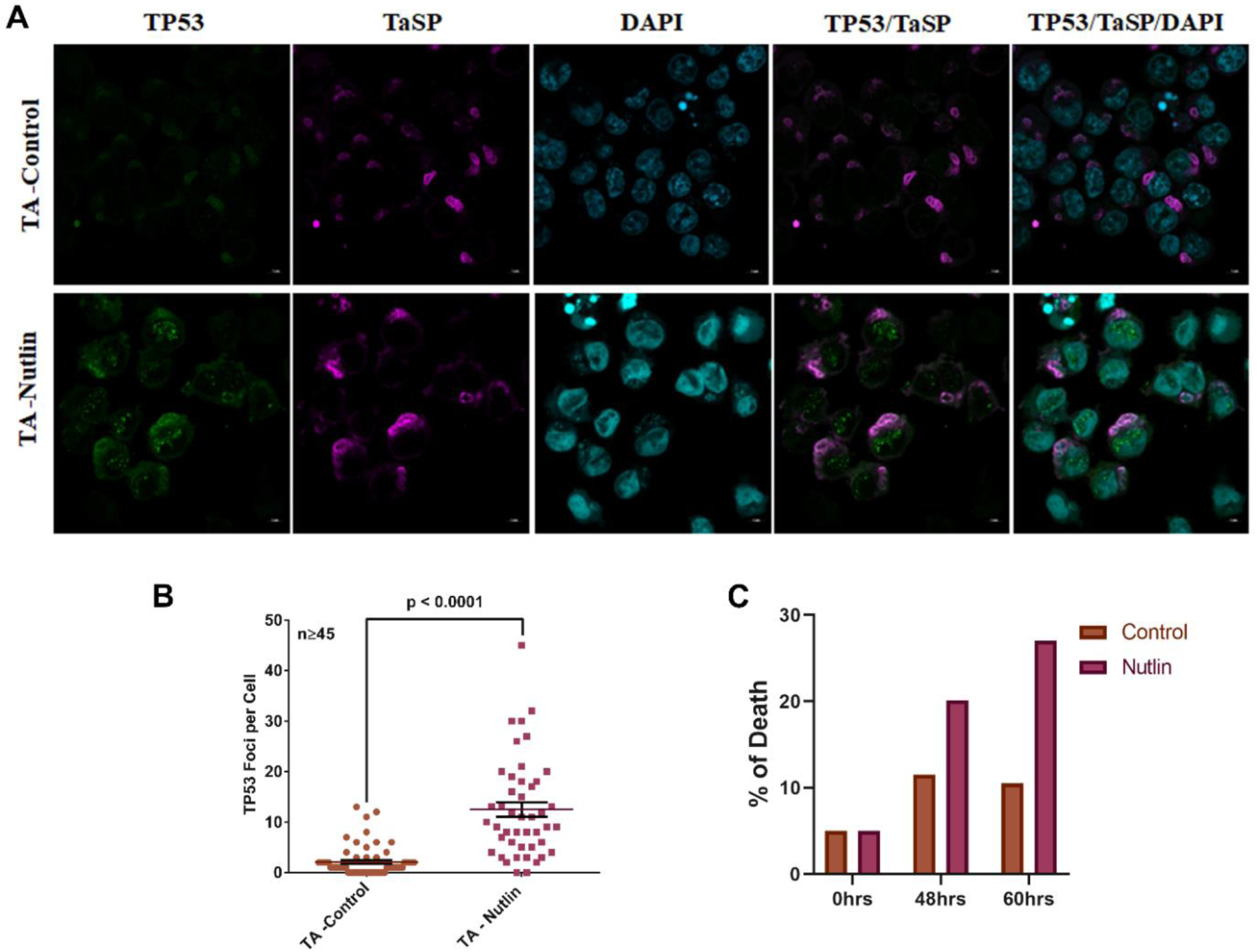
Effect of Nutlin-3A treatment on *T. annulata* infected bovine lymphocytes. A. Immunofluorescence analysis of *T. annulata* infected cells after 48 hrs treatment with Nutulin-3A using anti-p53 (Green) and TaSP antibody (Magenta). DAPI was used for nuclear staining. B. The graph demonstrates the activation of TP53 post Nutilin-3A treatment in *T. annulata* infected cells. C. The graph depicts the percentage of dead *T. annulata* infected cells post Nutilin-3A treatment at 0hrs, 48hrs and 60 hrs.

**Figure 8:**
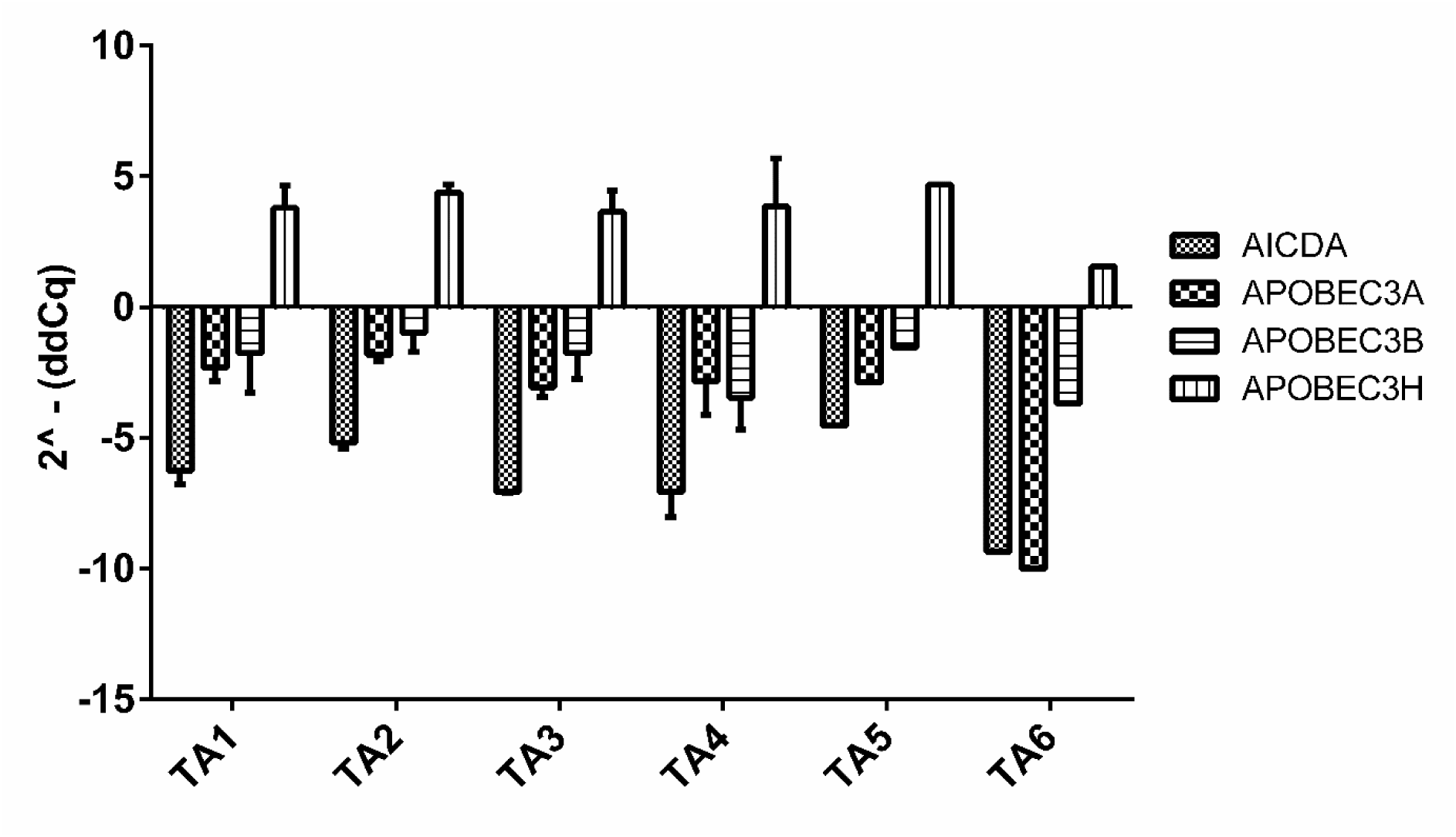
Expression levels of AID family genes in *T. annulata* infected cells: qPCR-based quantification of AID family genes in *T. annulata* infected cells in comparison to uninfected healthy PBMCs.

#### Cytidine Deaminase APOBEC3H might Contribute to Cancer like Mutagenesis in *Theileria-infected* cells

We next wanted to find the origin of somatic mutations, which we have discovered in our WGS analysis that might be helping the host cells to get immortalized. Recurrent infection with *Plasmodium* parasites has been shown to promote genomic instability and AID-Dependent B Cell Lymphoma (Robbiani *et al.,*2015). We next investigated the possibility of such events in the *Theileria-infected* cells by checking the expression of common DNA mutators like activation-induced cytidine deaminases (AID) family genes, specifically AICDA and APOBEC gene family members. We examined APOBEC3A, APOBEC3B, APOBEC3H and AICDA mRNA expression levels in our cell lines by qPCR. Within *Theileria*-infected cells, there was a substantial increase in APOPEC3H mRNA expression, while the expression of other genes exhibited a decline when compared to cells from healthy animals. These findings indicate that APOPEC3H could potentially contribute to somatic mutations in our cells.

## Discussion

This paper introduces the framework of somatic mutations observed in six clinically isolated T. annulata-infected leukocyte cell lines. These mutations have the potential to induce genomic instability in infected leukocytes, thereby contributing to their transformation resembling cancer-like characteristics. The WGS data were generated on infected leukocytes isolated from different animals harboring the same *T. annulata* parasite. This approach validly compares parasite-induced changes while removing variability caused by host genetic backgrounds and environmental exposure. To identify cancer-like somatic mutations, we compared the data with publicly accessible datasets from either healthy animals or the COSMIC and TCGA databases. As the goal was to identify genetic links between *Theileria*-transformed leukocytes and cancer we restricted our attention mainly to mutations in gene coding regions, as they have been extensively documented for a role in carcinogenesis.

The SNP analysis identified alterations in gene coding regions (n=7867) common to six *Theileria*-infected leukocyte cell lines. Comparative analysis using the TCGA and COSMIC databases allowed the identification of genes that are often changed in most cancers. The analysis revealed mutations in genes implicated in several cancer-related cellular and enzymatic processes, as shown by 127 distinct SMGs in *Theileria*-infected leukocytes. Comparison with the COSMIC database detected mutations in “ hallmark” genes known to be genetically associated with human cancer. The identified SNPS often did not occur at the same position, as those described in human cancer due to variations between human and bovine genomes. All the SNPs found in cancer-related genes are noteworthy and they might impact the malignant phenotype of *Theileria*-infected leukocytes. What was particularly striking was non-synonymous SNPs in 12 of the COSMIC tier-1 group signature genes. These signature genes might define the genetic predisposition of *Theileria*-infected bovine leukocytes to develop a cancer-like phenotype, because of their established role as oncogenes (FLT4, NOTCH2, MAP3K1, DAXX, FCGR2B, ROS1), tumor suppressor gene (TSG) (BARD1, KMT2C, GRIN2A, BAP1), or fusion genes (SLC34A2, NOTCH1) (Bamford *et al*., 2004). Among the 12 genes examined, homozygous mutations were discovered in GRIN2A, MAP3K1, KMT2C, and DAXX (Varmus 1984; Zhao *et al*., 2005; Bielski *et al*., 2018). TSG inactivation due to homozygous mutations or oncogene activation due to heterozygous mutations may contribute to the cancer-like features of *Theileria*-infected leukocytes. Loss of activity of GRIN2A, a well-known TSG, and activation of oncogenes (MAP3K1, DAXX, FLT4, NOTCH2, FCGR2B, ROS1) might contribute to *T. annulata*-induced transformation in infected leukocytes.

SNPs occurred in genes implicated in the genomic instability pathway (DAXX, BARD1, KMT2C, and BAP1), which is critical for cancer development, but has never been investigated in *Theileria*-induced leukocyte transformation. Genomic instability is a characteristic trait of nearly all cancer types and may be induced by endogenous (e.g., replication issues), or external (e.g., radiation) agents, and it’s almost always the consequence of deficient or abnormal DNA repair processes (Yoshioka *et al*., 2021; Li *et al*., 2020). SNPs were observed in DNA repair and replication pathway genes that are often mutated in cancer, and their presence increases the likelihood of host genome instability in *Theileria*-infected leukocytes. One of the essential DNA repair genes, CHEK2, harbored SNPs in all six *Theileria*-infected leukocyte cell lines. CHEK2 is a cell cycle checkpoint kinase that acts as a tumor suppressor, and defects in CHEK2 have been linked to an increased risk of cancer (Stolz *et al*., 2011). ROS1 had the highest number of SNPs in all six *Theileria*-infected leukocyte cell lines, so taking a hint from cancer research, we used crizotinib to inhibit the activity of bovine ROS1. Crizotinib inhibited the proliferation of *Theileria*-infected leukocytes that subsequently died. Overall, SNPs were identified in genes that should improve understanding of the cancer-like phenotype of *Theileria*-infected leukocytes and initiate the development of targeted therapies for the treatment of tropical theileriosis.

SNP signatures were extracted from the WGS datasets and compared to established cancer signatures to understand the basis of somatic mutation and genomic instability in *Theileria*-infected and transformed leukocytes. This uncovered two main cancer-like signatures indicative of a defective MMR and 5-methyl cytosine (5mC) deamination. MMR involves 5mC deamination-induced mismatches, and its absence in *Theileria*-infected leukocytes may result in an SBS1A mutational signature (Fang *et al*., 2021). Mutations in MMR pathway genes and polymerase proofreading enzymes (DNA Polymerase Epsilon, Catalytic Subunit (POLE) and DNA polymerase delta 1, catalytic subunit (POLD1)) are all linked to defects in DNA replication (Haradhvala *et al*., 2018). POLE and POLD1 genes that play a critical role in DNA replication and repair harbored multiple SNPs in all six *Theileria*-infected leukocyte cell lines. In cancer cells mutations in DNA repair genes underlie many forms of genomic instability, including MSI and chromosomal instability (CIN) (Negrini *et al*., 2010). MSI was observed in all *Theileria*-infected leukocyte lines indicating that parasite infection may generate host genomic instability, most likely due to defects in DNA mismatch repair activities. Mutations in the DAXX gene that can operate as either a TSG or an oncogene have been demonstrated to affect chromosomal stability and telomere maintenance in cancer (Heaphy *et al*., 2011; Lovejoy *et al*., 2012; Marinoni *et al*., 2014). DAXX mutations were consistently detected, and loss of DAXX protein and alternative telomere extension may be related to CIN in *Theileria*-infected leukocytes. In cancer, negative regulation of DAXX dampens the cellular apoptotic response and is linked to Pin1-mediated prolyl isomerization. Pin1 is a conserved pathway in *Theileria*-infected leukocytes to regulate host oncogenic signaling (Ryo *et al*., 2007; Marsolier *et al*., 2015).

We queried our dataset for CNVs and SVs, two forms of alteration that are known to play a significant role in cancer and influence a greater percentage of the genome than SNPs (Sudmant *et al*., 2015; Mahmoud *et al*., 2019; van Belzen *et al*., 2021). Ten cancer-related genes had copy number alterations in all six *T. annulata*-infected leukocyte cell lines. Amplification of specific loci can impact the expression of NFE2L2, BCL2L1, PAX8, PARP10, MDM4, or BPTF to promote the stimulation of infected leukocyte proliferation (Danovi *et al*., 2004; Guichard *et al*., 2012; Di Palma *et al*., 2013; Nicolae *et al*., 2014; Yin *et al*., 2016; Richart *et al*., 2016). The activation of transcription factors NRF2 (NFE2L2), PAX8 or BPTF may be just as crucial during the cellular transformation induced by *Theileria* infection, as the activation of other transcriptional regulators such as Myc (c-Myc), Nuclear Factor-kappa B (NF-кB), AP1 has been described (Chaussepied *et al*.,1998 ; Kinnaird *et al*., 2013). Differential expression of the NRF2, PAX and PARP gene families have been linked to *Theileria*-induced alterations in the infected host leukocyte (Kinnaird *et al*., 2013). We also noticed NFE2L2 (NRF2) and BCL2L1 overexpression in the six *Theileria*-infected leukocyte cell lines. Given their importance in cancer and their function as transcription factors and anti-apoptosis pathways, we hypothesize that these factors could potentially play a significant role in the immortalization of the bovine leukocytes. Furthermore, our study demonstrates that leukocytes infected by *T. annulata* exhibit aneuploidy-like cancer cells. This condition could potentially be triggered by copy number variations (CNV), or genome instability in infected leukocytes. Both aneuploidy and chromosomal instability have been widely acknowledged as fundamental characteristics of cancer (Hanahan D, Weinberg RA. 2000; Hanahan D, Weinberg RA. 2000). We hypothesize that copy number alterations are critical events in the evolution of the cancer phenotype in the *Theileria*-infected leukocytes, given the above observations of highly conserved CNVs across all six infected cell lines.

*Theileria*-infected leukocyte cell lines exhibit irreversible and reversible variations relating to changes in gene expression and epigenetic processes that contribute to developing a malignant phenotype <u>(Chaussepied *et al*.,1996; Tretina *et al*.,2015)</u> Like cancer, it is possible that genetic and epigenetic mechanisms are not independent events, but they interconnect and benefit each other to contribute to the cancer-like phenotypes of *Theileria*-infected leukocytes (You & Jones., 2012). One of the bovine genes found to be altered, KMT2C, is known to control the activity of DNMT3A a de novo DNA methyltransferase and silencing of its expression via histone methylation has been associated with cancer metastasis (You & Jones., 2012). Although we did not determine if the somatic alterations in key cancer-related genes affected epigenetic regulation, our results highlight the complexity of changes that could result from such alterations.

Research has indicated that more than half of human cancers have mutated TP53 genes in their somatic cells (Olivier *et al*., 2010; Rivlin *et al*., 2011; Muller & Vousden *et al*., 2013; Zhu *et al*., 2020; Solares and Deborah., 2022). However, our study did not reveal any mutations in this gene. This is in line with earlier work, which identified buparvaquone (BPQ) as a viable treatment for parasites, resulting in activation of TP53 and death of host leukocytes (Haller *et al*., 2010). Additionally, our discovery of Nutilin3a, which activates the TP53 gene and terminates the infected leukocytes, is in agreement with this. While SNPs have been identified in genes regulating TP53’s activity, such as TP53RK and TP53TG5, it remains to be seen if these SNPs affect TP53’s ability to induce leukocyte apoptosis. We hypothesize that Theileria infection inactivates TP53 by binding it to Mdm2, thereby permitting the host cells to become immortalized (Muller *et al*.,2013; Hafner *et al*.,2019). We believe it is the TP53 gene that plays a major role in immortalizing the host cells, not the parasite.

Pathogens such as Epstein-Barr virus (EBV), hepatitis C virus, HIV, Helicobacter pylori, and Plasmodium falciparum have been associated with a greater risk of B cell lymphoma and other cancers (Robbiani *et al*., 2015). Upregulated expression of the APOBEC3H gene suggests a resemblance to cancer, whereby this cytidine deaminase might contribute to the increased number of SNPs observed in *Theileria*-infected leukocytes. In addition, the APOBEC3 protein also plays a crucial role in the innate immune response to various viruses linked to the development of lymphomas, such as Epstein-Barr virus (EBV) and human T-cell leukemia virus (HTLV) (Wagener *et al*., 2015). Several studies have demonstrated the critical role of APOBEC3H, APOBEC3B, and APOBEC3A in the formation of mutations in numerous cancer types, like head/neck, lung, cervical, and bladder cancers (Starrett *et al*.,2016; Burns *et al*.,2013a; Burns *et al*.,2013b; Alexandrov *et al*., 2013; Zainal *et al*., 2012). However, further research is needed to understand better the connection and mechanism of the somatic mutations and cytidine deaminase related to *T. annulata*-induced bovine leukocyte transformation.

## Conclusion

We conducted an extensive study to investigate the occurrence of somatic mutagenesis throughout the bovine host genome in *Theileria*-infected leukocytes. This provided valuable insights into infection-induced degradation of host genome integrity and the mechanisms employed to counteract it, such as DNA repair processes. Our findings establish a solid foundation for unravelling the genetic events associated with the development of the cancer-like phenotype of bovine leukocytes infected by *T. annulata*. Notably, infection-induced overexpression of APOBECH3H has the potential to induce mutations in DNA repair genes, which may contribute to host genomic instability. Its elevated expression facilitates the accumulation of genetic alterations in DNA that likely contribute to the development of cancer-associated traits of *Theileria*-infected leukocytes. Although we detected SNPs in cancer-related genes, treatment with BPQ or Nutulin 3A causes apoptosis of the infected cell, indicating that interactions between host and parasite are vital for immortalizing the cell. We hypothesise that although cancer-related genes possess single nucleotide polymorphisms (SNPs), they cannot immortalise host cells unless TP53 is suppressed (Fig 9). Identifying SNPs in genes associated with DNA damage repair and genomic instability holds the potential to develop novel therapies targeting tropical theileriosis. This groundbreaking advancement will significantly enhance our fight against this disease.

**Figure 9:**
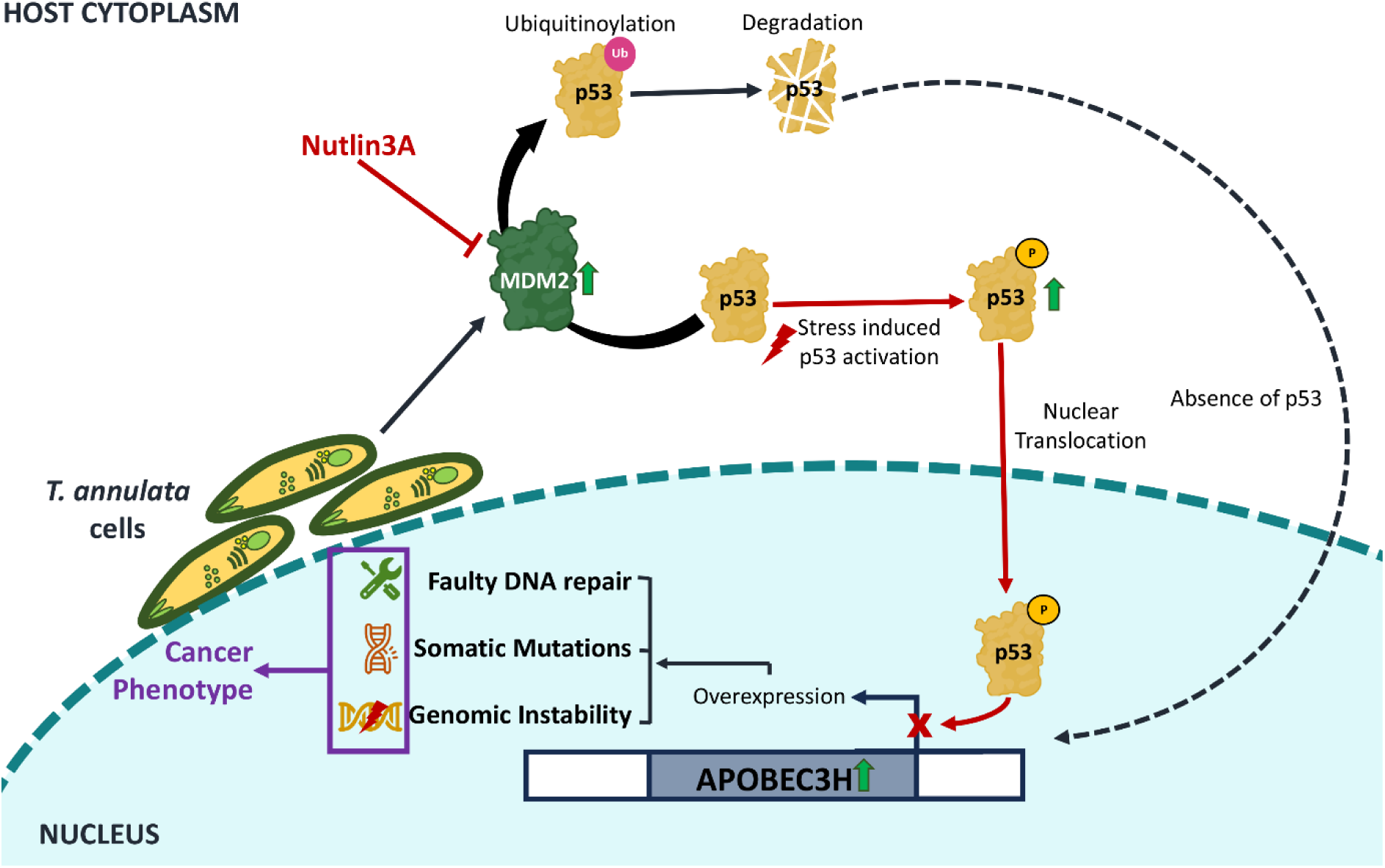
Proposed mechanism leading to p53 suppression, APOBECH3H upregulation and genomic instability in *T.annulata* infected cells.

## Materials and Methods

### DNA samples

The *T. annulata*-infected bovine cells (n=6) were isolated from the PBMCs of six infected animals. Cells were maintained in their standard medium to expand before DNA extraction. The DNA was extracted using QIAamp DNA Mini Kit (Qiagen) from the *in vitro T. annulata*-infected bovine cells, according to the manufacturer’s recommendations. Only isolated DNA with A260/280 ratios above 1.8 and proven high quality by gel electrophoresis were used for sequencing. All six samples were used for the WGS sequencing. Following the quality check, DNA samples were kept at −80 °C until further studies. Polymerase chain reaction (PCR) utilising primers specific for the *Theileria annulata* surface protein (TaSP) gene of *T. annulata* indicated the presence of parasites in host cells. The following are the PCR conditions for theaforementioned primers:

TaSP: 95 °C for 3 min, followed by 35 cycles of 95 °C for 1 min, 55 °C for 1 min, 72 °C and 1 min and a final extension of 5 min at 72 °C 5 min.

#### WGS analysis of *T. annulata*-infected bovine cells

The DNA from *T. annulata*-infected bovine cell lines TA1, TA2, TA3, TA4, TA5, and TA6 were submitted for sequencing with at least 30x coverage on the Illumina HiSeq sequencer (150X2 library type) using standard methods. We trimmed raw reads for adapter sequence removal, and reads with a minimum read length of 50bp, and minimum base quality Q30 were selected using the Trimgalore-v0.4.4 tool. The adapter free good quality reads were mapped to reference Bos taurus genome (Ensembl database: ARS-UCD1.2:CM008168.2) using Bowtie2-v2.0.5 at default parameters to generate alignment data (BAM format). The alignment data was processed using the Samtools-v1.9 program to generate mpileup data. We used Varscan2 (mpileup2snp and mpileup2indel) to predict variants (SNPs/indels) with various parameters such as minimum coverage (read depth) of 10, varfreq (variant frequency) of 0.2, phred base score of 30 (average quality score), and p-value threshold of 0.01 after filtering out insignificant and PCR duplicate reads (Koboldt *et al*., 2012). The predicted variants (SNPs and INDELs) were annotated based on the reference genome and gene feature information using the SnpEff-v3.3h tool. The predicted SNPs were reported as homozygous (1/1) if the variant allele frequency was a minimum of 60%. SnpEff and Ensembl Variant Effect Predictor (McLaren *et al*., 2016) were used to determine the effect of these variations on the genes (VEP). SnpEff annotates genomic variants and coding effects such as synonymous or non-synonymous amino acid substitution, start or stop codon gains or losses, or frameshifts (Cingolani *et al*., 2012). VEP employs the Sorting Intolerant From Tolerant (SIFT) method to forecast the impact of variations on the gene. SIFT spans from 0 to 1 and is based on a normalised chance of spotting the new amino acid at that place. A number between 0 and 0.05 is likely to impact protein function negatively. The WGS data generated in this study is submitted to the NCBI BioProject database (https://www.ncbi.nlm.nih.gov/bioproject/) under accession number PRJNA914920.

#### COSMIC and TCGA cancer database

COSMIC (Catalogue of Somatic Mutations in Cancer) is a publicly available database that collects and curates information on mutations in cancer (Tate *et al*., 2019). It contains data on a wide range of cancer types and includes information on the type and frequency of mutations, their location in specific genes and the functional consequences of these mutations. The Cancer Genome Atlas (TCGA) is another large-scale cancer database that provides comprehensive genomic information on various types of cancer (http://tcgaportal.org/). TCGA collects and analyzes data from thousands of cancer patients and provides information on gene expression, DNA methylation, and genetic mutations, among other data types. Both COSMIC and TCGA are valuable resources for researchers and clinicians studying the genetic basis of cancer or cancer-like diseases, as they provide large amounts of data that can be used to identify new therapeutic targets and to better understand the genetic changes that contribute to cancer development and progression. We compared our WGS variant-carrying gene list with COSMIC and TCGA gene lists. The information on genetic mutations in these databases was used for identifying actionable mutations in our samples.

#### Mutational signature and Microsatellite Instability

To find out the prevalence of any mutation type in the *T. annulata* infected samples we converted all mutations from the WGS data sets into a matrix composed of 96 single base substitutions for each mutation type (C > A, C > G, C > T, T > A, T > C and T > G) using each possible 5′ and 3′ contexts for all samples by applying the R package (SomaticSignatures) (Gehring *et al*., 2015). The distribution of mutational proportion for every 96 types are provided in the Supplementary Table (Supplemental_Table3.xls). After identifying the proportion of 96 mutational types we applied the R package deconstructSigs to infer the actual mutational signature (Rosenthal *et al*., 2016). In total, 30 signatures reported by COSMIC (http://cancer.sanger.ac.uk/cosmic/signatures) are included in the analysis. To further validate the signature of miss match repair deficient, we ran MSIsensor2® to identify the MSI level across *T. annulata* infected cells (Niu *et al*., 2014). As suggested by the literature, MSI scores greater than 20 were regarded as MSI, and values less than 20 were considered microsatellite stable.

#### Copy Number Variation and Structural variation

CNVs, or copy number variants, were identified using CNVcaller (Wang *et al*., 2017). We used 800 bp overlapping sliding windows to achieve this, and the results were compared across the samples. A population read depth file is created from the averaged read depths of all samples, and from this file, the potential CNV windows are selected. The potential CNV windows are merged into the CNV area if the distance between the two initial calls is less than 20% of their total length and the Pearson’s correlation index of the two CNVRs is significant at the 0 = 0.01 level (CNVR). Sample clustered read depth and individual integer copy numbers were used to identify the variant genotype. Two statistical analysis Silhouette score and the Calinski-Harabasze score, were used to find the significant CNVs regions. Structural variants were computed using Breakdancer to call for large inter and intra-chromosomal changes (Chen *et al*., 2009). The Breakdancer variant caller identified longer variants (>1KB), such as inversion, duplication, translocation, insertion, and deletion. Only structural variations with a confidence score of 99 were retained to maintain SVs with high confidence.

#### Real-time PCR

We performed Real-time PCR (q-PCR) by isolating RNA from *Theileria*-infected cells using the MN RNA isolation kit according to the manufacturer’s instructions. cDNA was synthesized from the RNA using reverse transcriptase (clonetech). cDNA isolated from the PBMCs of the healthy animal was used as a control for the q-PCR. For q-PCR, primers for the following ten genes were designed and synthesized. The sequences of the primers are listed in Table S? Using the Biorad machine’s default settings, we examined the relative expression of the target genes using HPRT as the reference gene. 2-ΔΔCT was calculated to evaluate the gene’s expression by comparing cDNA from healthy animals and *Theileria*-infected cells. Each experiment was conducted in triplicate.

#### Non-coding analysis

The variants not falling within any protein-coding genes were classified as non-coding variants. They are visualized in the Manhattan plot.

#### Cell culture and drug susceptibility assay

T.annulata infected cells were cultured in RPMI 1640 medium supplemented with 10% heat-inactivated fetal bovine serum at 37°C with 5% CO2. A challenge experiment with Crizotinib was carried out to assess the susceptibility of *Theileria*-infected cells to drug pressure (Cat no:12087; Cayman Chemicals). After 48 hr, each well received 20 µL of 1.5 mM resazurin dye, and the fluorescence intensity of the cells at 570 nm was measured 8 hr later to test cell survival. Untreated cells were used as a control. Likewise, host cell cytotoxicity experiments were performed on PBMCs collected from healthy cattle that tested negative for *T. annulata* infection. Each experiment was repeated three times.

#### Flow cytometry

DNA flow cytometry was utilized to evaluate the DNA content of *T. annulata*-infected cells in order to ascertain their ploidy status. The DNA staining process employed a propidium iodide-based method with BD flow cytometry. A standard reference sample was established by collecting blood from non-infected animals and subjecting it to DNA staining. The ploidy level was determined using the DNA index (DI), which divides the test sample’s mean fluorescence intensity (MFI) by the reference standard. A DI ratio greater than 1.1 and less than 1.9 indicates the presence of aneuploidy (Risques *et al*.,2001).

#### IFA

IFA experiments were done to verify the presence of T.annulata parasites and TP53 expression in the B-lymphocytes cells using a TASP antibody (1:200) raised against the *Theileria* annulata-surface protein and TP53 antibody(1:100). The IFA experiment and analysis were conducted following the methods described in a previous study [35]. In summary, 5 × 105 *T. annulata* infected cells were cultured and subsequently pelleted. The pelleted cells were washed three times with 1X PBS and then fixed with 4% paraformaldehyde at 37 °C for 10 minutes. Following fixation and permeabilization (0.1% Triton X-100), cells were then incubated in a blocking buffer (2% BSA in 1X PBS) at room temperature for 1 hour. Subsequently, the cells were exposed to primary antibodies overnight at 4 °C. After the washing step; the cells were incubated with a secondary antibody at room temperature for 1 hour. Following another round of washing, the DNA within the cells was labelled with DAPI. Finally, the samples were mounted using mounting media and images were captured using a fluorescent microscope. The acquired images were processed and quantified using the ZEN 3.3 (blue edition) software program.

#### Data access

Included in this paper are all datasets created and analyzed during this investigation. The whole-genome raw sequencing reads data generated in this study have been submitted to the NCBI BioProject database (https://www.ncbi.nlm.nih.gov/bioproject/) under accession number PRJNA914920. Additional data may be obtained from the appropriate author upon request.

## Acknowledgements

The authors would like to express their gratitude to the Director (NIAB) for providing all the necessary resources and for her consistent support and encouragement. We thank DBT JRF fellowship program (DB), CSIR JRF grant (AS and SS) and UGC JRF (MS) for providing fellowship to the PhD students. DD expresses gratitude to the Regional Centre for Biotechnology (RCB) in Faridabad, India, for allowing him to pursue his Ph.D. The results shown here are in whole or part based upon data generated by the TCGA Research Network: https://www.cancer.gov/tcga.

## Author contributions

PS and DD devised the study’s idea, planned the experiments, and evaluated and interpreted the results. Most of the experiments were carried out by DD. Data analysis was done by DD, RB, JG, AS, SS, MS, AT, VB and PS. The manuscript was written by DD, AS, SS, MS, VB, and PS. All authors were given a chance to discuss the findings and provide feedback on the article.

## Corresponding authors

Correspondence to Paresh Sharma.

## Transparency declarations

None to declare.

---

**Supplemental_Figure 1.pdf:**
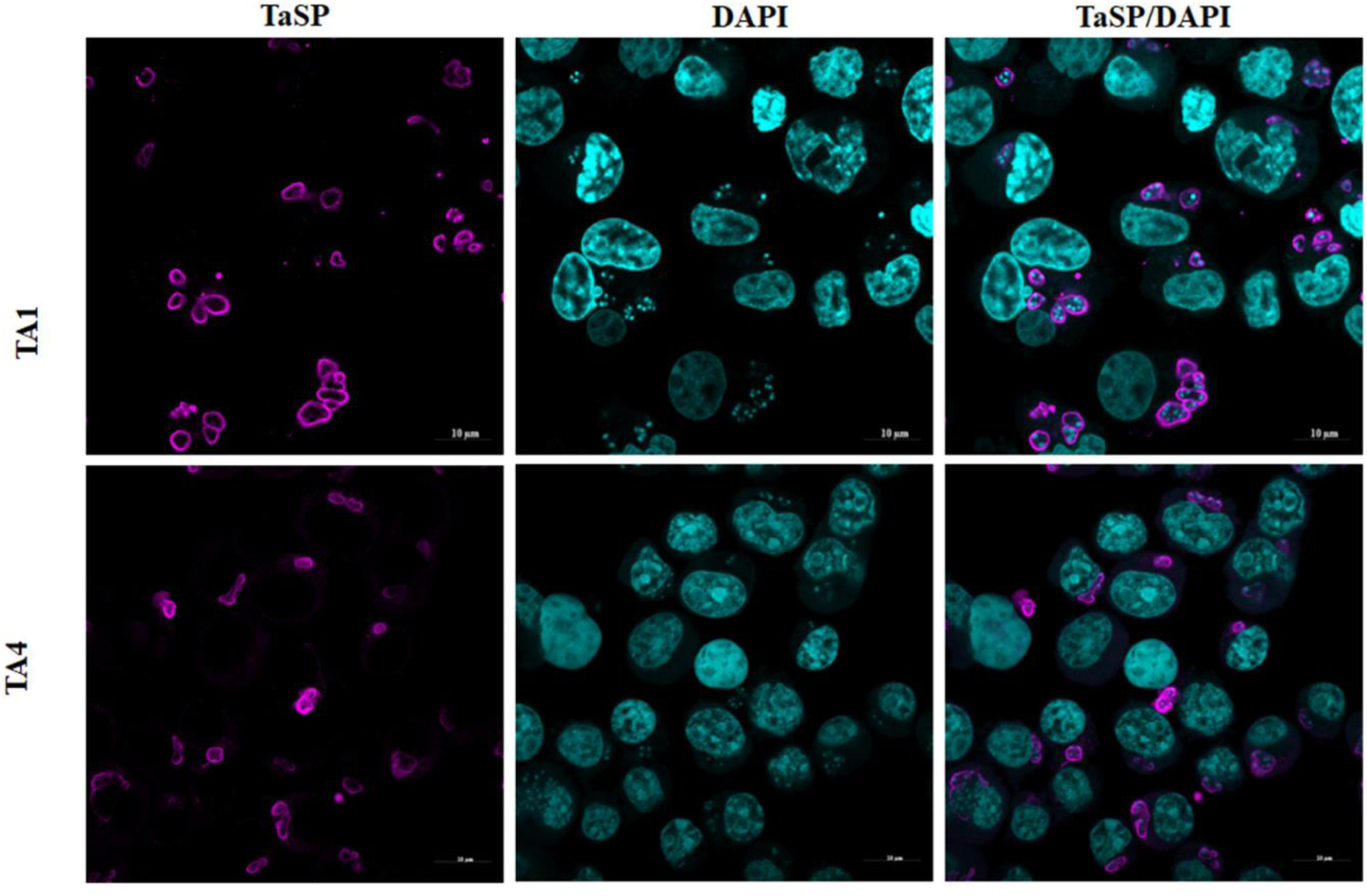
Immunofluorescence analysis of *T. annulata* infected cells, using TaSP antibody and DAPI was used for nuclear staining.

**Supplemental_Fig2.pdf :**
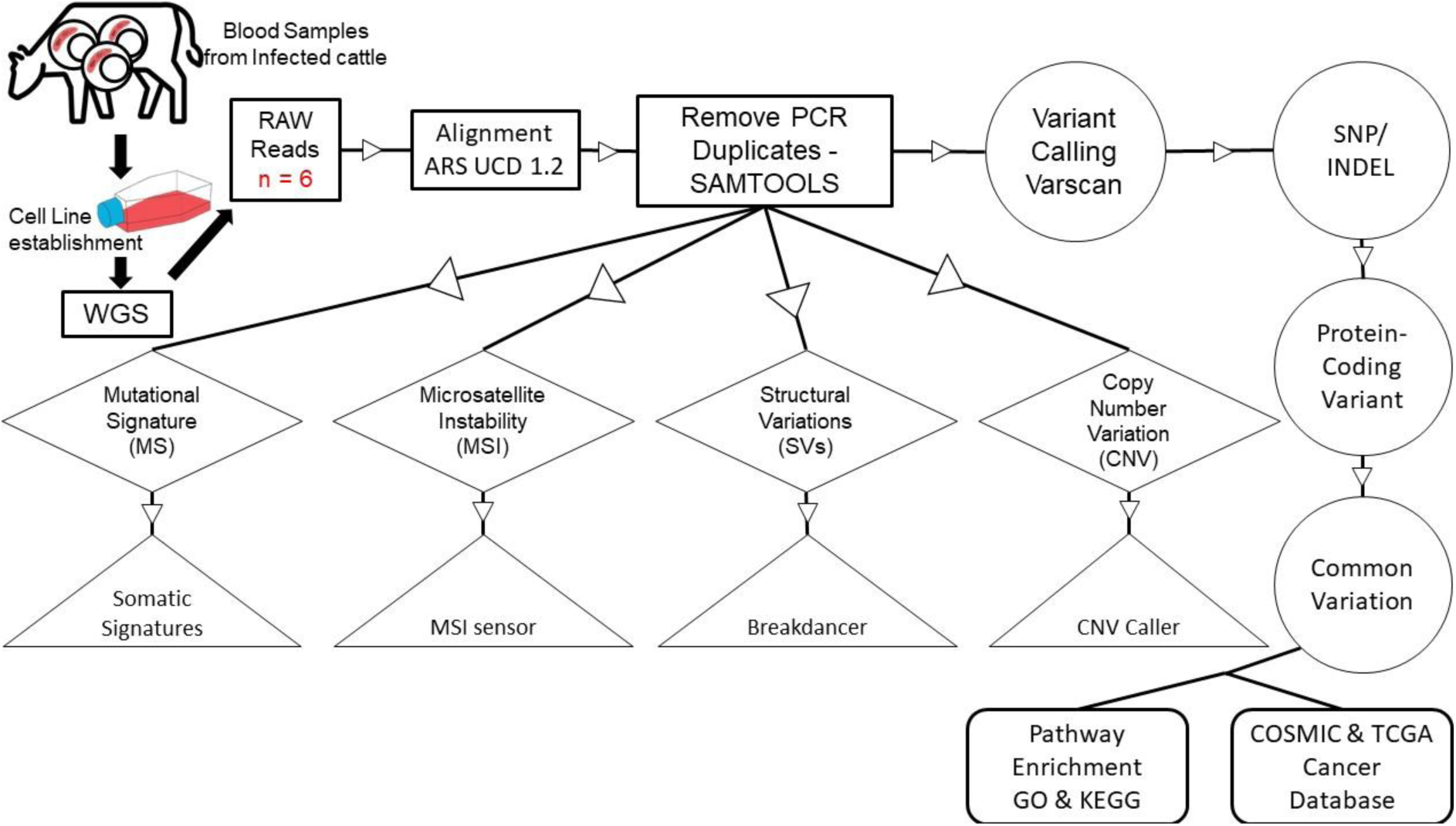
Schematic illustration of workflow: The process begins with the cell line establishment of *T .annulata* infected cells obtained from bovine blood samples of infected animals. 6 cell lines were established that were subjected to WGS. The RAW reads were aligned against ARS UCD 1.2 *B. taurus* genome. SamTools was used to process the aligned data. Further using Varscan SNPs and InDels were checked across all six samples with respect to the reference genome. The protein-coding common variants were selected and their significance in the development of a cancerous phenotype was assessed using COSMIC and TCGA databases. In parallel, from the aligned data the mutations in noncoding RNAs were also noted and were checked for various other genomic anomalies. These anomalies include MS, MSI, SVs, and CNVs which were calculated using the somatic signature, MSI sensor,Breakdancer, and CNV caller respectively.

